# Pre-trained Vision Transformers for Seizure Prediction: A Reproducible Baseline with Event-Based Evaluation and Statistical Validation

**DOI:** 10.64898/2026.03.11.711230

**Authors:** Ziyuan Yin, John Moraros, Shuihua Wang

## Abstract

**Background:** Scalp electroencephalography (EEG) based seizure prediction plays a critical role in improving the quality of life for patients with drug-resistant epilepsy, offering the potential for real-time warnings and timely interventions. Despite its clinical significance and decades of research, the field still lacks an open benchmark with reproducible baselines and deployment-oriented event-level evaluation. Most prior work relies on the small and outdated Children’s Hospital Boston (CHB-MIT) dataset and reports window-level metrics only, leaving the false-alarm burden of a real warning system underspecified. In seizure prediction, the cost of false alarm is significantly high since patients may receive painful electrical stimulation to suppress seizure. Hence, false alarms per hour (FA/h) and partial AUC (pAUC) are the most deployment-relevant metrics, reflecting alarm burden and discriminability in the low-false-alarm operating region that a usable warning system can realistically tolerate. However, few studies have systematically reported such metrics. In addition, vision transformers’ event-level performance under deployable FA/h constraints remains underexplored, and newer backbones such as MambaVision have yet to be evaluated under this setting.

**Methods:** In this work, we introduce a reproducible 5-fold benchmark derived from the Temple University Hospital EEG Seizure Corpus (TUSZ) dataset, and evaluate models using a pseudo-real-time event pipeline, reporting event-level sensitivity, false alarms per hour (FA/h) and partial AUC (pAUC). All models are compared to random predictors for statistical validation. We benchmark pre-trained vision transformers (SegFormer and MambaVision) under three EEG-to-image encoding methods, including a self-proposed Temporal-Patchify encoding for SegFormer.

**Results:** Our proposed Temporal-Patchify encoding method achieves state-of-the-art performance. We achieved 0.61 pAUC, which is 16.2% higher than the baseline Temporal-Tile SegFormer of Parani et al. The false-alarm burden (0.40±0.28 FA/h) is 44.4% lower than the Temporal-Tile SegFormer baseline while maintaining clinically usable sensitivity (60.7%±5.0%). We further perform statistical validation against a matched Poisson random predictor, confirming performance exceeds chance. Finally, we report end-to-end inference through-put up to 920 windows/s, confirming MambaVision’s fastest inference speed, exceeding SegFormer by over 20%.

**Conclusions:** This work bridges the gap between seizure prediction algorithms and clinically usable seizure prediction systems in real-world settings. Our findings indicate that pre-trained vision transformers, when coupled with appropriate EEG encoding methods, can achieve robust performance in low–false-alarm operating regimes, which is critical for real-world deployment. This benchmark and evaluation framework may facilitate more clinically meaningful and reproducible seizure prediction research.

## 1 Introduction

Epilepsy is a neurological disorder characterised by sudden, abnormal electrical discharges in the brain. Epileptic seizures typically result in muscle spasms, breathing difficulties, and loss of consciousness, significantly threatening patients’ safety and quality of life [1]. While most epilepsy patients can control/alleviate seizures with medication, approximately 30% of patients do not respond to medication [2] and are named as refractory epilepsy. For refractory epilepsy patients, invasive craniotomy is needed to remove the epileptic focus. But patients with generalised epilepsy (i.e., without focal lesions) or with difficult-to-localise lesions can not remove the epileptic focus by craniotomy [3].

The major challenge proposed by epilepsy is its unpredictability; seizures can occur suddenly and, without timely care, may in some cases lead to sudden unexpected death[4].

Therefore, a system capable of timely and accurate seizure prediction could significantly enhance safety and quality of life for these patients.

However, translating this clinical need into a practical warning system requires predicting seizures with an acceptable false-alarm rate under continuous EEG monitoring, a challenge that some researchers have attempted to address.

In 2013, J. Cook of the University of Melbourne, through long-term invasive EEG using NeuroVista, first demonstrated through clinical trials that epilepsy can be predicted via EEG [5]. A series of works based on Cook data, including the International Open Source Competition, have demonstrated the feasibility of predicting epilepsy through EEG [6]. However, these rigorously validated and open-source efforts primarily rely on invasive intracranial EEG, which is not suitable for routine daily use.

For real world, seizure prediction must be achieved with non-invasive scalp EEG. The transition to scalp EEG, while necessary, introduces significant challenges. Scalp EEG tend to be more noisy and dynamic.

Both scalp EEG and intracranial EEG collect changes in the pericerebral electric field caused by changes in the charge distribution of neurons in the brain. As scalp EEG and intracranial EEG originates from the same source, the two modalities exhibit consistent signal characteristics [7], and certain seizure events are reflected in both[8]. Motivated by this fact, many studies have attempted to use scalp EEG to predict epilepsy [9–11].

Early approaches attempted to extract nonlinear features from EEG [12] such as dynamical similarity index and effective correlation dimension [13]. To reduce reliance on manual feature design and better capture temporal structure, subsequent studies gradually adopted deep learning techniques, to predict epilepsy from handcrafted features by using LSTM and RNNs [6, 14]. More recently, transformers and their variants, including vision transformers, have been investigated for EEG analysis because of their capacity to model long-range dependencies and learn more expressive latent representations [15].

Despite methodological advances and improved performence, the field still lacks the systematic assessment of models’ performance on the latest public dataset and an open and reproducible benchmark.

The datasets commonly used in scalp EEG seizure prediction are typically small in size and lack population representativeness.

Most studies only report prediction results based on a single public dataset, CHBMIT, released 16 years ago (2009), which comprises only 22 paediatric patients. Paediatric brains are still developing and differ substantially from those of adults. Consequently, the onset of most neurological diseases in children differs substantially from that in adults [16]. Moreover, in most studies using CHB-MIT data, the average number of seizures per patient is only 9, resulting only a few seizure events used for test[17, 18].

Despite many studies claim near-perfect accuracy, these inflated results are often attributable to unintentional data leakage. Another critical issue is that many studies confuse sample-level detection with event-level detection. As illustrated in Fig. 1, one epileptic seizure is considered a single event, and each event corresponds to a batch of warning zone samples cut out according to a specific Seizure Prediction Horizon (SPH) and Seizure Occurrence Period (SOP). The smallest unit input to the seizure prediction model is a sample, and the predicted probability output by the model is also based on a sample. Therefore, all samples originating from the same seizure event must be kept entirely within one set—training, validation, or testing—to ensure independence. However, some studies unintentionally combine all event samples for random partitioning, thereby unintentionally improving the metrics and achieving the so-called 100% prediction. The lack of open benchmarks and reproducible baselines has worsened this problem, causing inflated good but not verified results.

**Fig. 1.**
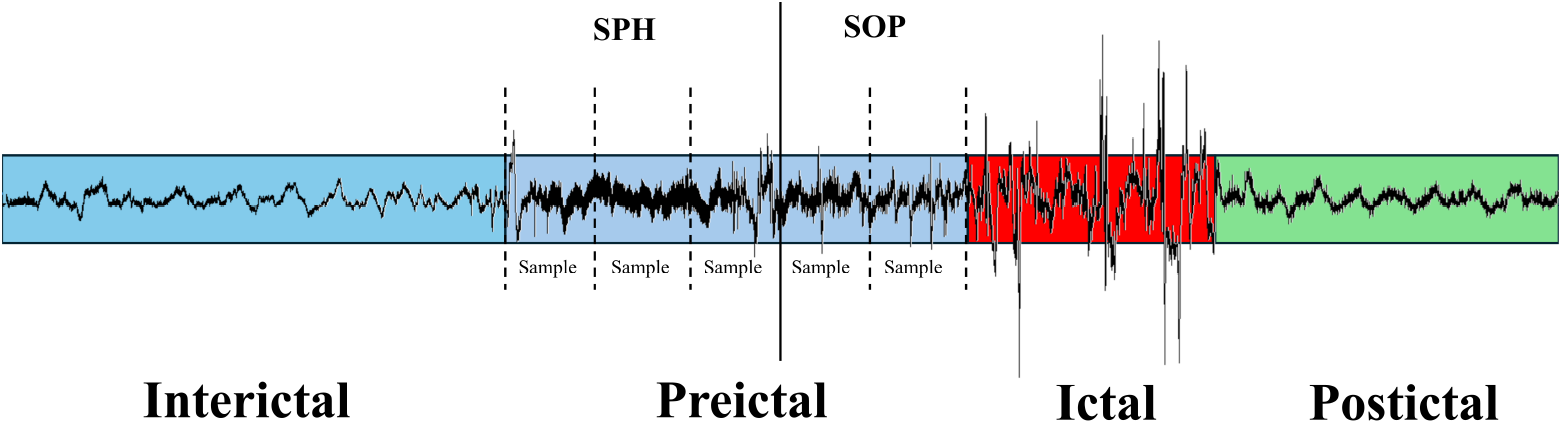
The four distinct states of the EEG signal of an epilepsy patient.

Moreover, event-level metrics of the models are overlooked. Current research mainly assesses metrics at the sample level, i.e., the model’s ability to classify segments as positive or not, rather than its performance on predicting whole seizure events. This oversight hinders the investigation into whether the current model is clinically ready or not. Several patient interviews suggest that analysing epilepsy system metrics from an event-level perspective is crucial [19]. For example, excessively high time-in-warning (TIW) values can cause patient anxiety, rendering the device ineffective in clinical practice. A low false-positive rate is extremely critical in real-world applications: numerous studies indicate that excessively high false positives substantially weaken user trust, lead to device abandonment, and are among the core obstacles preventing current prediction algorithms from being adopted in clinical settings. Many methods lead to a discrepancy where the system “looks accurate but is actually unusable. Such metrics can only be calculated at the event level.

Hence, determining the clinical utility of an algorithm requires prospective or pseudo-prospective evaluation of sensitivity and measurement of false alarm rate (FA/h) [20]. However, current models simply compare sample-level ROC-AUC metrics, which provide limited clinical insights without considering FA/h. A system with high ROC-AUC but high FA/h would be impractical, as the cost of triggering a false alarm is significant and cannot be tolerated [21]. Additionally, the traditional ROC-AUC metric does not work and cannot reflect a system’s event-level performance since the area under the curve, but right to the vertical FA/h cut-off line, is meaningless. Any operating point with FA/h over a specified value disables the system.

Even with event-level metrics, one fundamental question remains: under the same false-alarm budget, is the observed sensitivity genuinely better than what could be achieved by chance? All systems should be compared with random predictors for statistical validation [2], a step that has been overlooked in past research.

To bridge these research gaps, this paper proposes a fully open-source, reproducible benchmark derived from the largest public epilepsy seizure dataset TUSZ. We establish a baseline for epilepsy early warning using two vision transformers as backbones with different EEG-image mapping methods. Model performance is evaluated at the event level for sensitivity, FA/h and partial AUC. The model is also compared with random guessing to demonstrate its statistical effectiveness. Results show that our proposed Temporal-Patchify Encoding mapping method outperforms prior SegFormer-based methods. Specifically, our contributions in this paper are as follows:

1. We propose a fully open-source, reproducible benchmark derived from the Temple University epilepsy seizure dataset containing 675 patients with 4029 seizures. The TUSZ dataset is much more representative than the CHB-MIT dataset, addressing the problem of poor generalisation caused by the past reliance on the small-scale CHB-MIT dataset.
2. We report event-level metrics of seizure prediction models on a publicly available dataset, including Sensitivity, FA/h and partial AUC, thus laying a baseline for future research. Real-world deployment readiness can only be evaluated based on these metrics.
3. To the best of our knowledge, We are the first to propose using the pre-trained lightweight MambaVision structure for seizure prediction. MambaVision enhances the speed of the seizure prediction model by over 20% by reducing the time complexity of the attention mechanism.
4. We propose a Temporal-Patchify Encoding module which transforms the raw EEG signal into images for input into SegFormer, and this module makes the model achieve the SOTA performance in terms of event-level metrics.
5. We systematically compare the model with random predictors, statistically demonstrating the model’s effectiveness through p-values greatly lower than 0.05.

## 2 Literature Review

Most seizure prediction studies follow two main paradigms: (i) classical feature-engineering pipelines (spectral/time–frequency and nonlinear dynamics descriptors) coupled with conventional classifiers, and (ii) end-to-end deep representation learning approaches (e.g., CNN/RNN/Transformer-based models) that learn features directly from EEG. In addition, statistical models of seizure timing provide complementary baselines for judging whether observed performance exceeds chance.

### 2.1 Early Feature-based and Statistical Approaches

Although Hans Berger first recorded human electroencephalography (EEG) in 1924 [22], systematic exploration for seizure prediction did not begin until the 1990s due to limitations in analytical tools. Early work focused on spectral and time–frequency analysis: for example, [23] used spectral estimation of dual-channel scalp EEG signals, combined with features such as coherence and pole trajectories, to achieve seizure prediction; Schiff et al. proposed the “brain chirps” model in the time-frequency representation of implanted EEGs and used matched filtering to construct a prototype for early signs [24]. In parallel, seizures were also modelled as temporal point events; statistical modelling treated seizures as a sequence of time events. Milton et al. showed that the seizure intervals of about half of the patients did not follow a Poisson distribution, suggesting temporal dependence and clustering characteristics within seizure EEG [25].

### 2.2 Nonlinear Dynamics and Feature Engineering

Meanwhile, researchers began to adopt nonlinear time-series analysis to capture dynamical changes preceding seizures. For example, Elger and Lehnertz [26] used non-linear time series analysis to show that the pre-ictal EEG exhibits a transition to a lower-dimensional state; Iasemidis et al. [27] used nonlinear indicators such as phase space topography and Lyapunov exponent to characterize the dynamic changes before ictal events; subsequently, Le Van Quyen [28] applied nonlinear similarity measures to clinical EEG and verified its real-time usability. Concurrently, engineered intelligent algorithms combine feature engineering with machine learning: Geva and Kerem [29] used wavelet features and unsupervised fuzzy clustering for widespread ictal prediction; D’Alessandro et al. [30] used genetic algorithms for multi-channel, multi-feature selection, combined with probabilistic neural networks, to construct an individual-oriented classification framework with a prediction window of approximately 10 minutes. In 2002, the IWSP1 conference in Bonn was the first to systematically bring together and align the aforementioned scattered research, unifying the topics and data practices, and forming a phased consensus for the clinical neurophysiology community.

### 2.3 Rigorous Evaluation Reveals Limitations

However, these early results were quickly tested under more rigorous evaluation: Aschenbrenner-Scheibe et al. [31] systematically evaluated nonlinear methods such as “dimensionality reduction” on long-term implantable EEGs, pointing out their under-performance under rigorous hold-out tests; Mormann et al.’s 2007 review [12] further emphasized that “the current literature allows no definite conclusion as to whether seizures are predictable by prospective algorithms.” These mixed findings highlighted the need for long-term recordings and rigorous prospective-style validation to establish whether prediction truly exceeds chance.

### 2.4 Breakthroughs with Invasive EEG and Deep Learning

Cook [5] pioneered seizure prediction using long-term implanted iEEG signals, providing statistically validated evidence of “superiority over randomness” and validating the feasibility of prediction, laying the foundation for subsequent research. However, the experimental algorithm used in this research was initially developed in 2008. Due to limitations in statistical feature extraction algorithms at the time, such as power spectral density, the seizure prediction results were poor for three patients. Subsequently, Kuhlmann [6] successfully solved the seizure prediction problem for patients with poor prediction results using deep learning, achieving the best ROC-AUC 0.81.

### 2.5 Deep Learning Advances: from CNN to Transformer

Following these breakthroughs, machine learning techniques, including CNN, LSTM, and Transformer, were gradually applied to seizure prediction. Truong et al. [32] used convolutional neural networks (CNN) to learn distinguishable signal features from STFT time-frequency spectrograms, achieving epilepsy seizure prediction in patient-specific scenarios involving iEEG and sEEG. Acharya et al.[33] used a one-dimensional CNN to directly extract features from raw EEG, promoting the application of end-to-end models in seizure prediction. Meng [34] proposes M-NIG (Mobile Network Information Gain), a correlation-network-based EEG seizure prediction framework that builds patient-specific dynamic connectivity graphs to improve noise robustness and highlight critical (DNB) channels, reporting improved sensitivity and reduced false alarms on CHB-MIT and a self-collected dataset. Tsiouris et al. [35] used LSTM to model long-term dependencies within the extracted features, improving patient-specific prediction performance; Wu et al. [17] further provided an end-to-end LSTM prediction process without manual features. Jina et al [36] combined a topological convolutional neural network with LSTM to model complex interactions between different brain regions, achieving cross-dataset generalizability.

More recently, transformer-based architectures have gained traction. Li [37] improved the transformer with a CNN structure, which overcomes the drawback of the transformer that it cannot extract local features exactly. Meanwhile, graph neural networks were used to model the spatial relationships [38] between EEG channels, to lighten the model and improve its generalisation performance. Zhu et al. [39] fused a Transformer and an RNN to capture the internal dependencies between long-term EEG signals and integrate multi-channel information. Qi [40] used an improved ViT based on a visual transformer for seizure prediction, while Parani [15] proposed a novel wrangling method to directly input the original signal into ViT for prediction and achieved sample-level ROC-AUC 0.87. Yuan et al. [41] used a hybrid architecture of DenseNet and ViT, utilising local convolution and global attention mechanisms to achieve complementary advantages.

### 2.6 Research Gap

Despite rapid progress in model design, comparisons remain difficult because many studies rely on small or outdated datasets, inconsistent splitting practices, and predominantly sample-level reporting. Deployment-oriented event-level evaluation under constrained false-alarm rates, together with statistical validation against chance predictors, is still underreported. These gaps motivate the event-disjoint TUSZ benchmark and protocol introduced next.

## 3 Dataset and Benchmark

This section introduces the TUSZ dataset and the method of processing TUSZ to form a benchmark for seizure prediction.

### 3.1 TUSZ Dataset Overview

The Temple University Hospital EEG Seizure Corpus (TUSZ) was established by the Neural Engineering Data Consortium (NEDC) to address the lack of a public dataset, despite the widespread collection of EEG data worldwide [42]. The data is sourced from the medical records database of Temple University’s affiliated research hospital, with the initial release comprising 14 years of historical records. Currently, Temple University’s Institute for Signal and Information Processing maintains and updates TUSZ, which offers greater diversity in seizure types, a larger patient cohort, and more transparent, standardised data collection and annotation processes than comparable datasets. TUSZ represents the latest generation of public datasets for deep learning applications, as summarised in Table 1. Its large, open-source nature makes it particularly suitable for training deep learning models.

**Table 1.** Comparison of representative public EEG datasets for seizure-related research.

| Time Period | Dataset | Patients | Hours | Seizures |
| --- | --- | --- | --- | --- |
| 1990s | Bonn EEG | 10 | 3.28 | 100 |
| 1990s | Freiburg iEEG | 21 | 509 | 88 |
| 2010s | CHB-MIT | 22 | 844 | 198 |
| 2010s | EPILEPSIAE | 275 | 40000 | 2662 |
| 2015–Present | TUSZ | 675 | 1476 | 4029 |
| 2018s | Siena Database | 14 | 128 | 47 |
| 2018s | Helsinki Neonatal | 79 | 97 | 460 |
| 2020s | SeizeIT2 | 125 | 11000 | 883 |

To enhance data usability and the interpretability of machine learning models, the data annotation standards are detailed in [43], including criteria for distinguishing between spike and sharp waves, ictal events, and artefact events. The latest dataset is TUSZ V2.0.3. It contains EEG data from 675 patients, 1645 sessions, 7377 EDF files, totalling 1476 hours, covering various types of epileptic seizures from focal to generalised epilepsy, and 4029 seizure events in total.

The TUSZ data structure is shown in Fig. 2. TUSZ has three file formats: edf (European Data Format) stores EEG signals, including sampling rate, acquisition date, and channel information. The channels (17 channels at least) were from the 10-20 international system, so each channel corresponds to a brain region that can be mapped using the 10-20 system. Most sessions employed a sampling rate of 256 Hz. However, some sessions utilised sampling rates of 250 Hz, 400 Hz, 512 Hz, or 1000 Hz. The csvbi file is a binary labelling file used to label whether a specific segment of the EEG signal indicates epilepsy, including duration, montage type, labels, and confidence level. The csvbi file contains the same content, but the csv file is a multi-class label, indicating the specific type of epileptic seizure, such as focal nonspecific seizure or generalised seizure. For the TUSZ dataset, there are 13 labels in total, as shown in Table 2.

**Table 2.** The labels used to annotate TUSZ data. Definitions adapted from [43].

| Labels | Meaning |
| --- | --- |
| bckg | Cerebral signals that do not correspond to seizure activity. |
| seiz | Seizure. General annotation indicating the presence of seizure activity. |
| fnsz | Focal nonspecific seizure. A broad category of seizures localized to a specific region of the brain. |
| gnsz | Generalized seizure. A broad category of seizures involving most or all regions of the brain. |
| spsz | Simple partial seizure. Brief seizures originating in a single brain region, during which the patient remains fully aware and responsive. |
| cpsz | Complex partial seizure. Similar to a simple partial seizure, but characterised by impaired awareness. |
| absz | Absence seizure. A brief, sudden seizure characterised by a lapse in attention, typically lasting less than 5 seconds and most commonly observed in children. |
| tnsz | Tonic seizure. A seizure involving the stiffening of the muscles. Usually associated with and annotated as tonic-clonic seizures, but not always (rarely there is no clonic phase). |
| cnsz | Clonic seizure. A seizure involving sustained, rhythmic jerking, typically observed in conjunction with tonic-clonic seizures and annotated accordingly. |
| tcsz | Tonic-clonic seizure. A seizure characterized by loss of consciousness and violent muscle contractions. |
| atsz | Atonic seizure. A seizure involving sudden loss of muscle tone, often occurring as a phase preceding a tonic-clonic seizure. |
| mysz | Myoclonic seizure. A seizure characterized by brief, involuntary muscle twitching or myoclonus. |
| nesz | Non-epileptic seizure. An observed event resembling a seizure but lacking electrographic evidence of epileptic activity. |

**Fig. 2.**
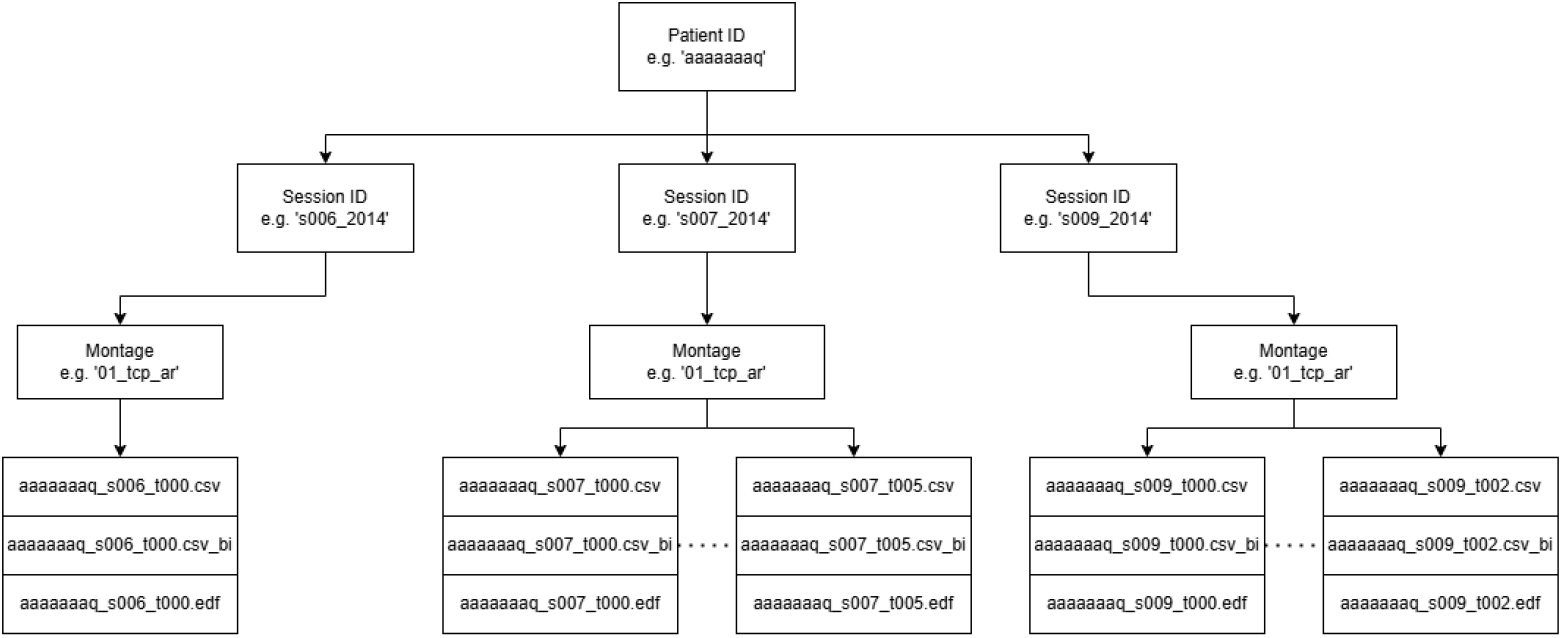
Top (e.g., aaaaaaaq) is the patient’s anonymous unique identifier. Each patient has a folder named after this ID. Under the patient folder is a ‘session’ folder, and each ‘session’ folder contains data from a specific session. For example, ‘s006_2014’ indicates that this folder contains data from session 006 collected in 2014 for this patient. Each ‘session’ folder contains a unique folder indicating the montage configuration. ‘tcp’ indicates the electrode placement is temporary central parasagittal, and ‘ar’ indicates the reference electrode is referenced to an average reference. ‘01_tcp_ar’ indicates that this session uses the ‘01 tcp ar montage’ configuration. For specific montage information, see [44]. The final level folder contains ‘csv’, ‘csv_bi’, and ‘edf’ files, representing multi-class annotation, binary annotation, and EEG data, respectively. Note that most of the data from 2002 to 2019 was collected using equipment from Natus Medical Incorporated. Natus uses a proprietary, confidential data format and will remove EEG files containing no information, as instructed by technicians [44]. Therefore, the data for each session may not be continuous, and some sessions’ EEG data may consist of multiple EDF files. Each file name includes a token, such as t000. In most cases, the tokens are arranged in chronological order, meaning that t002 will not be earlier than t001 [44], but the specific token order is determined by the technicians operating Natus.

Due to licensing restrictions, the source data cannot be distributed directly; researchers seeking access must apply to the NEDC for permission.

### 3.2 Benchmark

Electroencephalogram (EEG) recordings of epileptic patients are typically categorised into four distinct states [45]: interictal, preictal, ictal, and postictal, as illustrated in Fig. 1. The interictal state denotes the normal EEG activity observed during daily life, whereas the preictal state refers to the latent period preceding an epileptic seizure. Evidence shows that certain patients display specific EEG characteristics during the preictal period [46, 47]. The ictal state corresponds to the seizure event, characterised by substantial abnormal electrical discharges in the brain. The postictal state represents the recovery phase following a seizure, during which the chaotic high-frequency EEG peaks diminish, and the amplitude of the EEG signal envelope progressively decreases.

The TUSZ dataset was originally used to train epilepsy monitoring models, not prediction models. Therefore, it only annotates the ictal period of epileptic patients’ EEG signals; it lacks information about the preictal period. To convert TUSZ into a publicly available dataset usable for seizure prediction models, it is necessary to define the seizure prediction horizon (SPH), the time before an epileptic seizure during which the epilepsy system should issue an alert.

An ideal epilepsy warning system should be able to predict the exact seizure onset time, since current systems cannot predict the exact seizure onset time, the seizure occurrence period (SOP) is introduced to quantify the uncertainty window [20]. For a correct prediction, the seizure prediction system issues an alert within the SPH region, and the seizure occurs after the SOP interval.

The SPH time provides sufficient response time for epilepsy suppression systems, such as electrical stimulation or drug administration, ensuring effective seizure control. The SOP time allows sufficient time for the aforementioned suppression systems to take effect; however, it should not be excessively long, as a prolonged SOP time would increase side effects. In this study, both SPH and SOP were set to 5 minutes.

Under the condition of SOP = 5 min and SPH = 5 min, the conversion of TUSZ data for epilepsy monitoring involves the following steps. First, the sampling rates of the EEG signals stored in all EDF files need to be unified to 256Hz. Then, redundant information is removed from each EDF file, retaining only 17 channel signals, as shown in Table 3. Next, the EEG signal segments from each patient’s session are concatenated. As shown in Fig. 2, the EEG signals from each session are divided into several EDF files, and the temporal order of the EDF files is determined by the token.

**Table 3.** The original 17 channels preserved from the TUSZ dataset. Electrode labels follow the international 10–20 system; each entry indicates the electrode name and its approximate scalp region (L/R: left/right hemisphere; Z: midline).

| Channels | Channels |
| --- | --- |
| T6: R post-temp | FP1: L frontopolar |
| T5: L post-temp | F8: R ant-temp |
| T4: R mid-temp | F7: L ant-temp |
| T3: L mid-temp | F4: R frontal |
| P4: R parietal | F3: L frontal |
| P3: L parietal | CZ: mid central |
| O2: R occipital | C4: R central |
| O1: L occipital | C3: L central |
| FP2: R frontopolar |  |

Finally, positive samples for seizures that occurred more than SOP + SPH = 10 minutes after the previous seizure are extracted. Simultaneously, all patient sessions are traversed. If a patient has a session with no seizures, a 5-minute segment of the EEG signal is randomly truncated from that session, and negative samples are extracted using the same segmentation method. The specific algorithm flow is shown in Algorithm 1.

After successfully extracting samples for a seizure, the corresponding samples for that seizure are stored in the same.npy file, named patient session seizure token. That is, each.npy file stores the positive and negative samples corresponding to a specific seizure in a specific session of a specific patient. This storage method effectively prevents the data leakage risk mentioned in the introduction.

The dataset is then divided into five folds for cross-validation. This division is based on epileptic seizures, meaning that samples corresponding to a single seizure will only appear in either the training or validation set. However, samples corresponding to different seizures from the same patient are allowed to appear in both the training and validation sets.

This setup is based on the understanding that each epileptic seizure is an independent event, and the samples from the preictal period of a single seizure exhibit similar characteristics. Including samples from the same seizure in both the training and validation sets would lead to data leakage and inflate the model’s performance. The inclusion of samples from different seizures of the same patient across sets is justified because numerous medical studies, including those using invasive EEG, have confirmed [48, 49] that the preictal EEG patterns of each epileptic patient are not consistent, and there is currently no universal EEG biomarker.

Following this process, the TUSZ dataset is transformed into a 5-fold cross-validation dataset for epileptic seizure prediction, with each fold based on individual epileptic seizures. With the event-disjoint benchmark and preprocessing pipeline defined, we next describe the vision backbones and EEG-to-vision encoding strategies evaluated under this protocol.

#### Algorithm 1

Preictal and interictal samples extraction

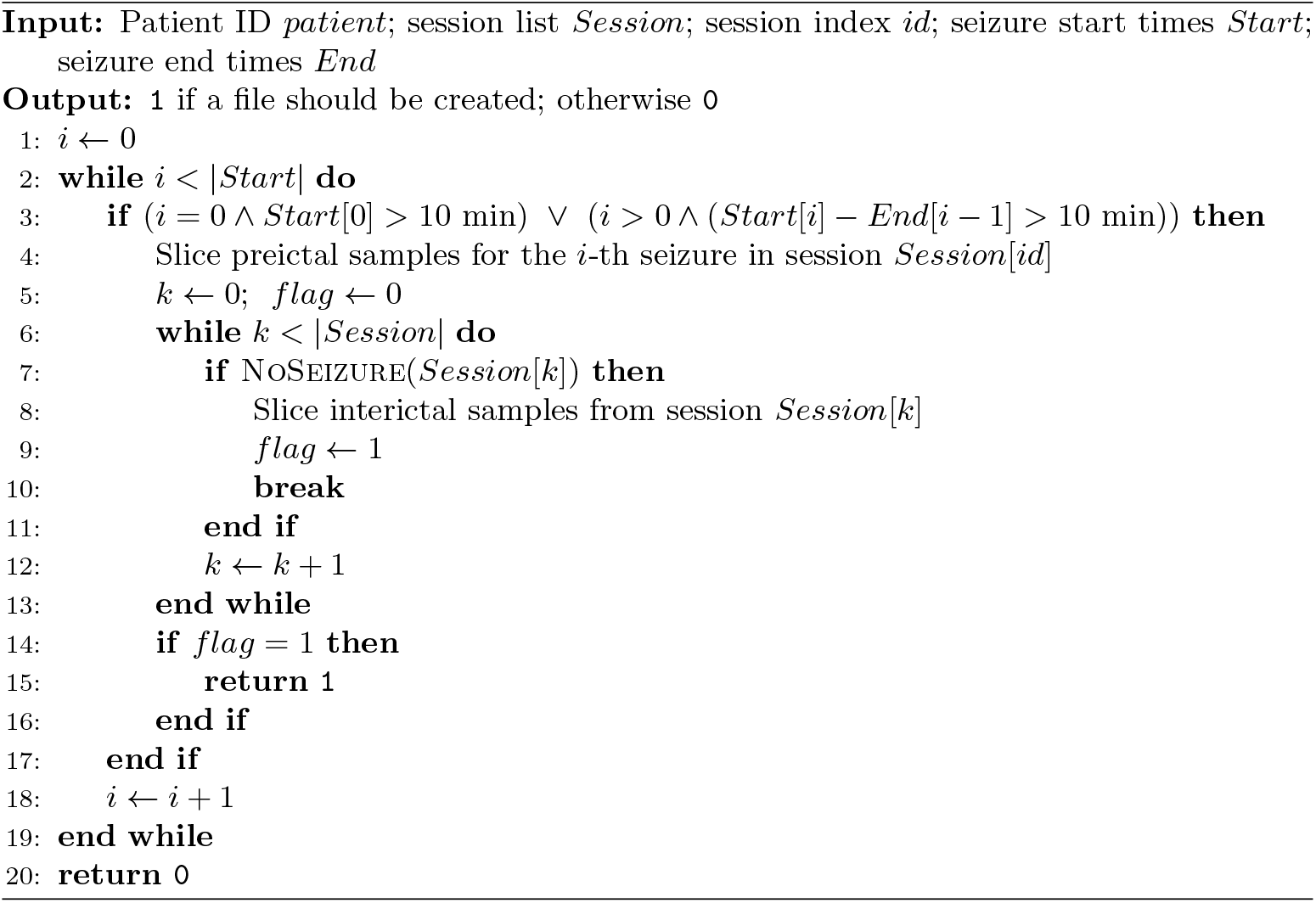

## 4 Model Architectures and Experimental Comparisons

To assess whether pre-trained vision backbones can serve as practical seizure forecasters under our event-level protocol, we evaluate two families of lightweight models—SegFormer and MambaVision—paired with three EEG-to-vision encoding strategies: SegFormer is a traditional vision transformer with a global attention mechanism, whereas MambaVision utilises a space-state machine to replace the attention mechanism for lower complexity. Different input encoding methods are proposed to transform the raw temporal EEG into representations suitable for models. This subsection introduces the two baseline model architectures. Then, implementation details and protocols are disclosed. Finally, the post-processing of sample-level predictions to event-level predictions and the design of a random predictor for statistical validation are presented.

### 4.1 Backbone Design

#### 4.1.1 Backbone-1

The first vision backbone is utilised from [50], namely SegFormer, which comprises 4 repeated stages: an overlap patch embedding, a transformer block, and a stage normalisation. The overall structure of the SegFormer is shown in Fig. 3. Each transformer block in each stage has an encoder and decoder with complexity *O*(*N*^2^) as illustrated in equation 1 below, where *Q, K, V ∈* ℝ^*N×d*head^ denote the query/key/value projections, *N* is the token (sequence) length, and *d*_head_ is the per-head embedding dimension.

**Fig. 3.**
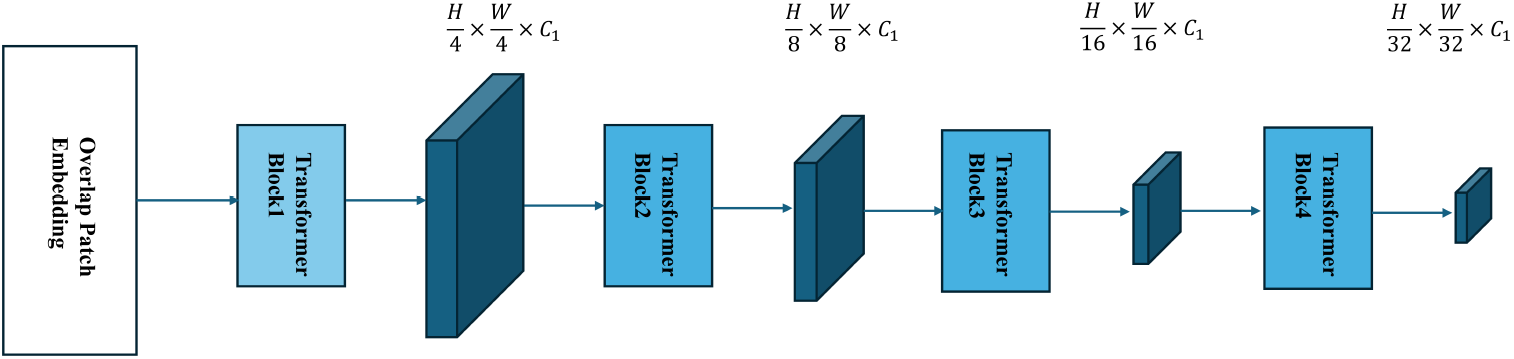
Illustration of SegFormer encoder structure.

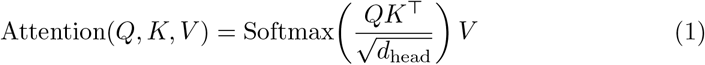

This would be prohibitive for large image resolutions [50]. The SegFormer employed a sequence-reduction process to shorten the sequence length, as shown in equations 2 and 3, where *N* is the token length and *C* is the embedding dimension. The reduction process is controlled by a reduction ratio *R. X ∈* ℝ^*N ×C*^ is the input token sequence, 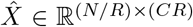 *∈* ℝ^(*N/R*)*×*(*CR*)^ is the reshaped sequence for reduction, and *X*^*′*^*∈* ℝ^(*N/R*)*×C*^ is the projected reduced sequence. The complexity of the attention mechanism is reduced from *O*(*N*^2^) to *O*(*N*^2^*/R*), and *R* is set to be 64,16,4 and 1, respectively, in each stage. In this research, the complexity would be reduced to *O*(*N*^2^*/*64), *O*(*N*^2^*/*16),*O*(*N*^2^*/*4) and *O*(*N*^2^) as a result.

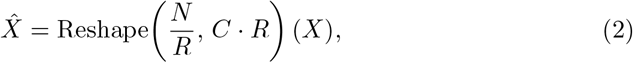

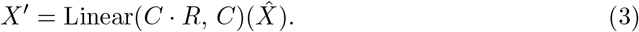

The encoder model utilised is from the NVIDIA/MIT-B0 checkpoint, which is fine-tuned on ImageNet-1k.

#### 4.1.2 Backbone-2

The second vision backbone is based on MambaVision [51], which includes a state machine and an attention mechanism, compared to 4 repeated attention blocks in the vision backbone-1. The complexity of MambaVision would be *O*(*N*) + *O*(*N*^2^). The overall structure is illustrated in Fig. 4.

**Fig. 4.**
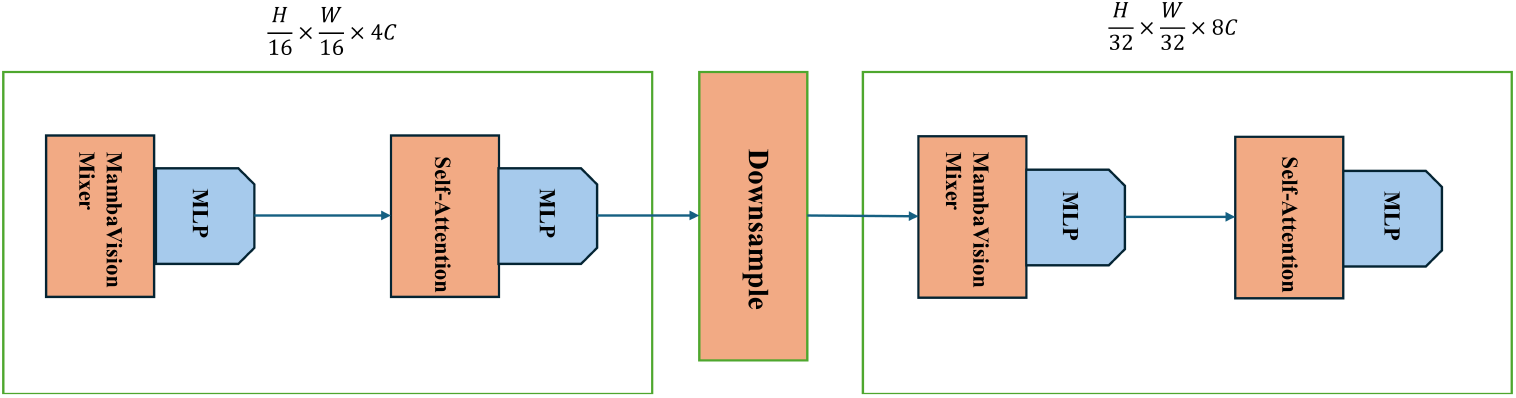
Illustration of MambaVision’s core structure.

The core of MambaVision is a block added before the attention block and is redesigned based on the original Mamba mixer [51]. The causal convolution is replaced with regular convolution to make it suitable for vision inference. Moreover, to compensate for content loss during the space state machine stage, a parallel branch without a space state machine is added, consisting of a convolutional layer and a sigmoid linear unit. The outputs of both branches are concatenated using a final linear layer.

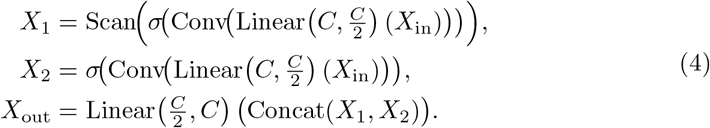

In equation 4, *X*_*in*_ is the input into the mixer and *X*_*out*_ is the output. *Linear* is the linear layer with the first parameter as the input and the second parameter as the output embedding. *σ* is the activation of the sigmoid linear unit mentioned above. *Conv* is the regular convolutional layer replacing causal convolution, *Concat* concatenates outputs of dual branches, *C* is the embedding dimension and Scan denotes the (selective) state-space scan operator. The output size of each branch is reduced to *C/*2. According to [51], such a dual-branch design enriches models’ capacity to extract high-level features for vision tasks.

The model utilised is from the NVIDIA/MambaVision-T-1K checkpoint, which is also fine-tuned on ImageNet-1k as vision backbone-1.

### 4.2 EEG-to-Vision Input Encoding

Following prior work by [52], the Temporal-Tile encoding with SegFormer is adopted as a baseline throughout this benchmark, serving as a reference point for evaluating alternative encoding strategies and models.

#### 4.2.1 Encoding-1 Temporal-Tile Encoding

To make time-series data suitable for vision backbones pre-trained on 512*512 images with 3 channels, three encoding methods have been proposed to convert (17, 512)- dimensional temporal EEG signals into images suitable for vision backbones. One of which is from [52], where EEG segments are replicated along the channel dimension to form a 512*512 image.

For any input signal *x*(*t*) with (17, 512) size *x*(*t*) goes through a linear mapping layer to be expanded into a size of (20, 512) signal *y*(*t*). Then replicate *y*(*t*) along its channel dimension 25 times to form an interim signal of size (500, 512). Finally, the first 12 channels of *y*(*t*) are selected and concatenated into the bottom of the interim signal to form *Y*, which is of size (512, 512) as illustrated in equation 5, where *x*(*t*) *∈* ℝ^17*×*512^ is a 2-s EEG segment, *y*(*t*) *∈* ℝ^20*×*512^ is the linearly mapped signal, *y*(*t*)_(*j*)_ denotes the *j*-th replication (tile), and *y*_1:12_(*t*) denotes the first 12 channels.

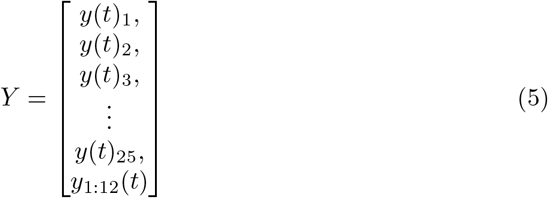

After being reshaped into (512, 512), the *Y* is then repeated three times to represent (R, G, B) channels respectively.

#### 4.2.2 Encoding-2 Temporal-Patchify Encoding

The second input encoding method first linearly remaps the input signal and then applies the process called patchify, proposed in this paper. This method aims to capture the tiny, high-frequency changes over a short period and enhance them by tiling. The overall structure is illustrated in Fig. 5.

**Fig. 5.**
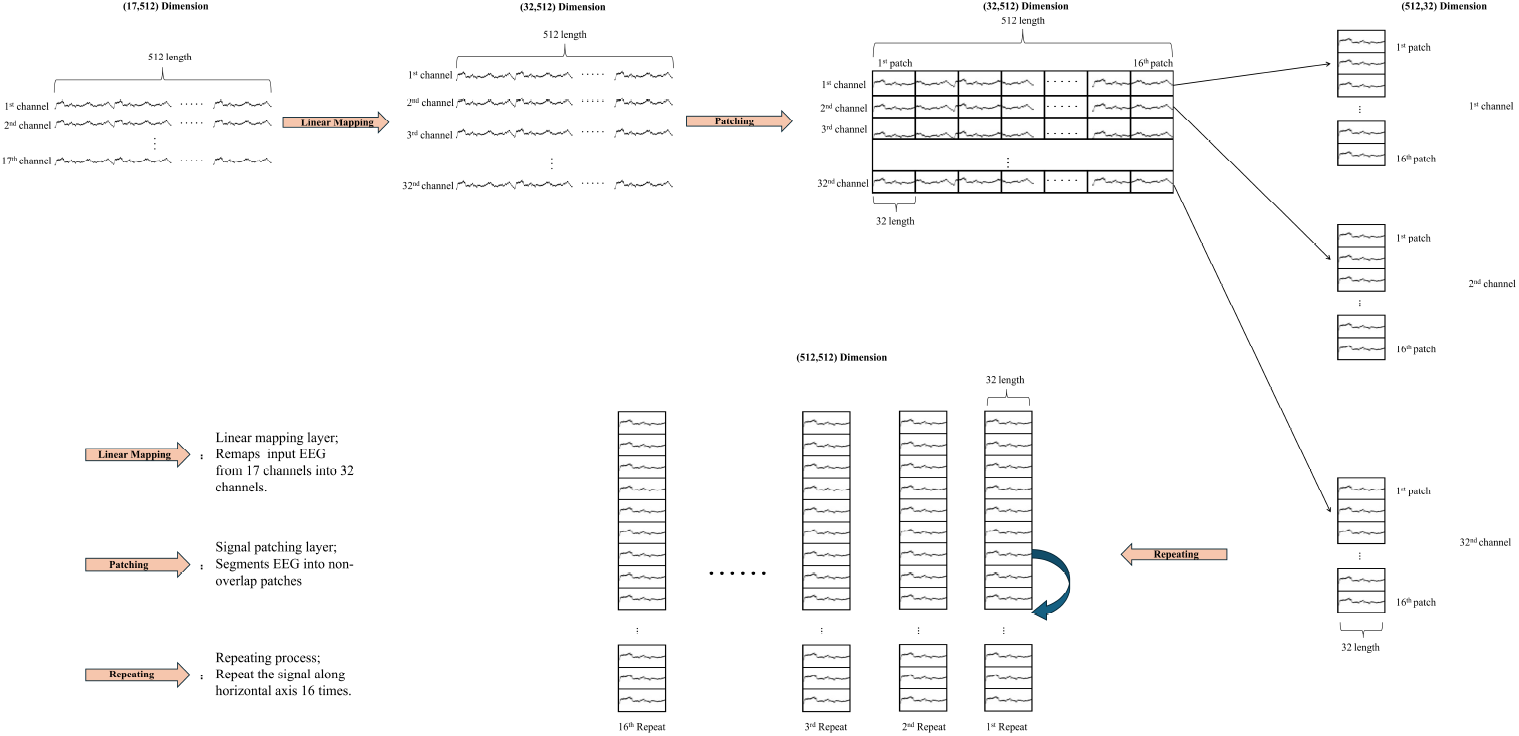
Demonstration of workflow of the Temporal-Patchify Encoding for transforming raw 17-channel EEG into a suitable input.

The input signal was first transformed into *Y* of size (32, 512) as shown in equation 6.

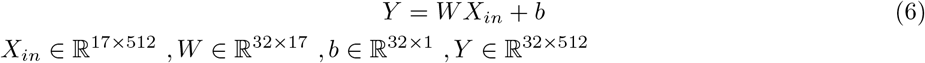

The detailed structure of *Y* is illustrated in equation 7 for the better understanding of Temporal-Patchify Encoding.

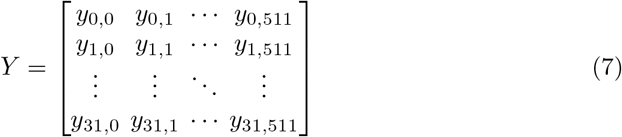

For each channel signal **y**_*i*,:_ *i ∈* (0, 31) with size of (1, 512), reshapes each channel into size of (16, 32) by tiling every 32 segments into *Y tiled*_*i*_ as shown in equation 8 where *y*_*i,a*:*b*_ denotes the contiguous samples from index *a* to *b*.

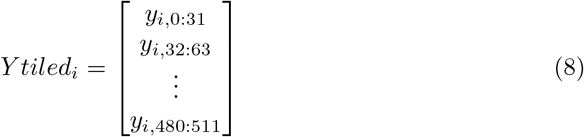

After transforming each channel into *Y tiled*_*i*_, concatenate each 32 *Y tiled*_*i*_ along the vertical axis to form a signal of size (512, 32) as shown in equation 9 where *Y*_tiled_ ∈ ℝ^512*×*32^.

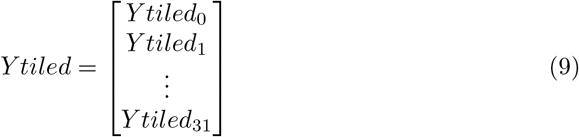

Finally, repeat *Y tiled* 16 times along the horizontal axis to form the signal size of (512, 512) as shown in equation 10 where Output *∈* ℝ^512*×*512^.

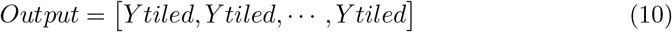

As in the first encoding method, the final signal is stacked 3 times to represent the (R, G, B) channels.

#### 4.2.3 Encoding-3 Linear-Mapping

The third encoding method is merely the linear mapping of raw EEG segments into different channel dimensions. This design could preserve the raw temporal information while leveraging a strong pre-trained vision backbone for detailed information extraction. This process mimics the diagnostic process of medical experts who review cross-channel information for seizure detection. Because raw 17 channels are signals collected from 17 separate electrodes, neurologists tend to manually check cross-channel information, such as the (Fp1-F7) channel, to visualise brain activity within that range. Instead of predefining which cross-channels to use, this research uses a linear mapping layer to learn which cross-channel information is suitable.

### 4.3 Event-Level Evaluation

As discussed in the introduction, there is a lack of evaluation of models’ performance on the event level. Real-time deployment metrics needs to calculate the sensitivity based on the number of correctly predicted seizure events rather than samples, as illustrated in the equation 11 where *N*_pred_ is the number of seizure events with at least one valid alarm and *N*_total_ is the total number of seizure events.

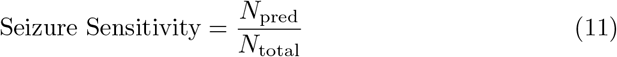

To calculate the above seizure sensitivity, the workflow is to first smooth a series of discrete predicted samples, then trigger an alarm if certain criteria are met. A refractory period is activated after an alarm is triggered.

In this research, for each seizure event with 2-second samples and SPH 5 minutes, a 150-length predicted sample series *p*[*t*], *t ∈ range*(0, 149) is filtered by a causal exponential moving average filter (EMA) as shown in the equations 12 and 13, where *α* is the smoothing factor, *t* is the sample index and 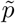 [*t*] denotes the EMA-smoothed prediction sequence..

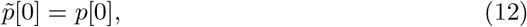

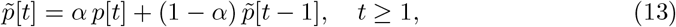

The exponential moving average introduces an effective temporal integration on the order of 1*/α* samples (approximately 5 seconds in our setting), which is substantially shorter than the adopted SPH, as shown in equation 14.

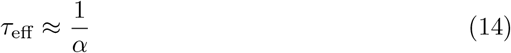

Finally, by adopting the K-out-of-N strategy, once *p*[*t*] exceeds the threshold consecutively in K samples, an alarm is triggered at *t*_*alarm*_. As long as *t*_*alarm*_ *∈* [*t*_*seizure*_ *− SOP − SPH, t*_*seizure*_ *− SOP*], it would be a correct prediction, where *t* is the sample index within a 5-min segment (*t* = 0, …, 149), *t*_seizure_ is the seizure onset time, and *t*_alarm_ is the time of the alarm.

In this research, the *α* of the EMA filter is set as 0.4, and K is set as 15. And the threshold is determined with different values which will be discussed in the results part.

### 4.4 Statistical Validation

For a prediction model to be effective, it must be shown to outperform any random or periodic predictor. Because for any given SPH, if SPH is long enough, any random predictor may trigger an alarm corresponding to a seizure event. A prediction model cannot be clinically ready if it achieves lower sensitivity than a random predictor under the same constraint.

To design a comparable random predictor, the false alarm rate (FA/h) has to be consistent with the prediction model, ensuring an identical false alarm burden between the random and proposed predictors. If FA/h is lower than that of the random prediction model, then the random predictor inevitably inflates sensitivities; vice versa.

The Poisson alarm model is adopted as the random predictor. Equation 15 shows the model where *N* (*t*) represents the number of alarms triggered during the time interval [0, *t*]. *λ* is the constant equal to FA/h.

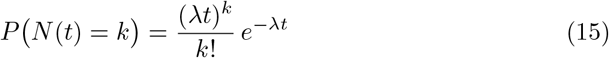

The model’s expectation is shown below, where *t* is measured in hours.

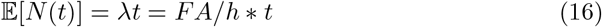

Hence, for any given period, the probability of having at least one alarm is shown in equation 17.

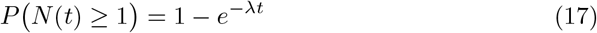

The sensitivity of a random predictor is the probability of having at least one alarm in the SPH interval, as shown below, where *T*_*SPH*_ is the SPH duration.

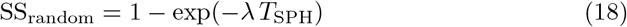

This SS_random_ will be compared with the sensitivity of the real predictor on the event level.

## 5 Results and Discussion

This section evaluates the proposed models using multiple metrics, including ROC-AUC, seizure event sensitivity, false alarm burden (FA/h), partial AUC, and inference latency. In addition, we compare all models against a random Poisson predictor to justify statistical effectiveness.

The results show that our proposed encoding method, Temporal Patchify, achieves SOTA partial AUC and enables the model to be deployable. Also, MambaVision, which is applied to seizure prediction for the first time, achieves the fastest inference speed.

This research explores four models in total, summarised in Table 4. The two different vision backbones introduced earlier are combined with different input encodings to transform the EEG signal into a form suitable for each backbone.

**Table 4.** Model variants with different input encodings and vision backbones.

| ID | Input encoding | Vision backbone |
| --- | --- | --- |
| 1 | Temporal-Tile | SegFormer |
| 2 | Temporal-Patchify | SegFormer |
| 3 | Linear mapping to (20, 512) | MambaVision |
| 4 | Linear mapping to (256, 512) | MambaVision |

**Table 5.**
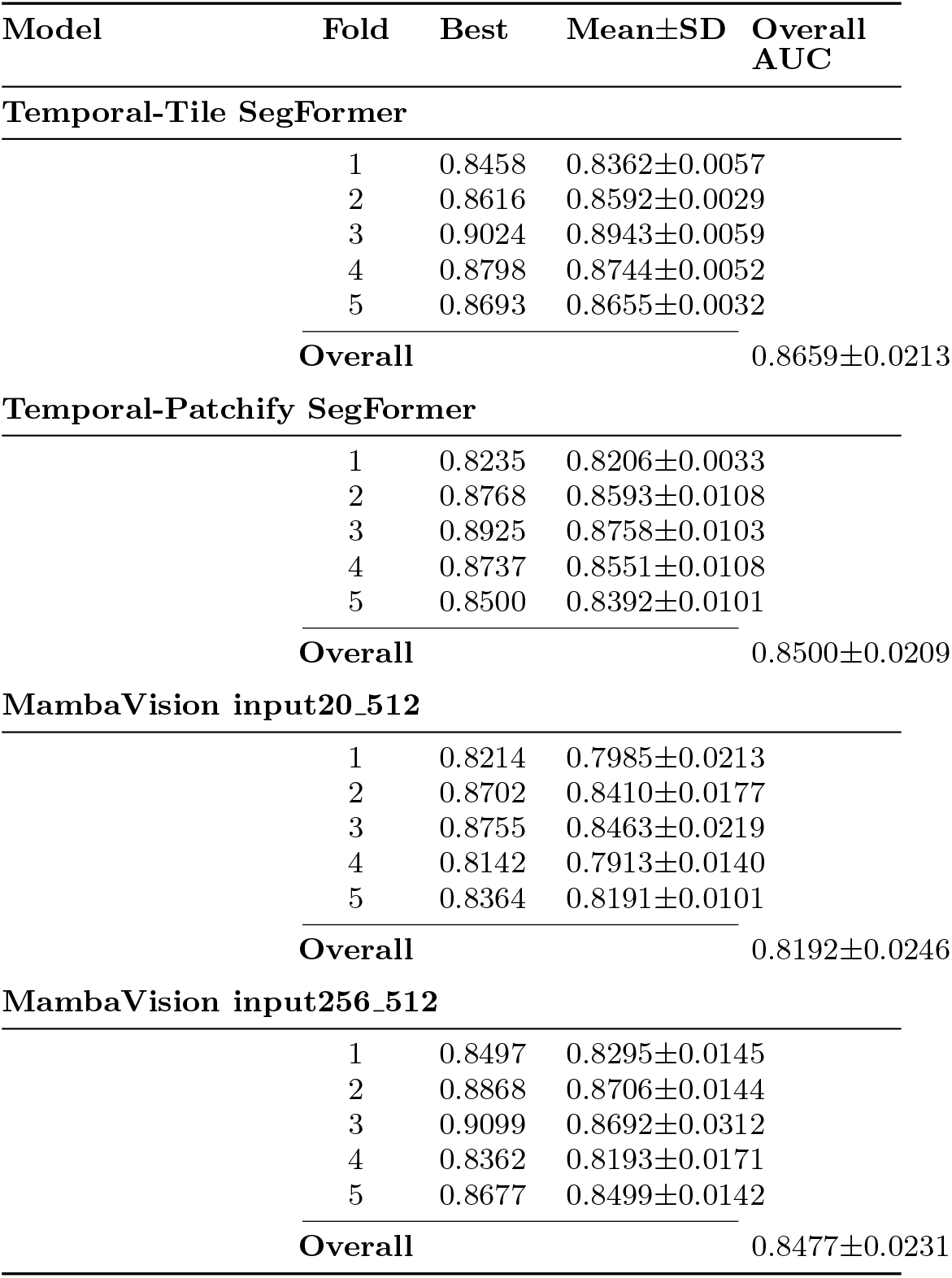
Fold-wise window-level ROC-AUC. “Best” is the maximum ROC-AUC among six runs within each fold; Mean ±SD are computed across the six runs. Overall AUC is Mean ±SD across the five folds using per-fold means.

### 5.1 Window-level ROC-AUC

Each model is trained and validated on all 5 folds with 6 runs in each fold, resulting in 30 runs for each model. Considering that the threshold affects models’ sensitivity, where a low threshold leads to high sensitivity, reporting sensitivity alone is insufficient for assessing a model’s ability to distinguish between interictal and preictal segments. Hence, this research reports ROC-AUC to assess its performance on the sample level. The summary is presented in Table 5.

#### 5.1.1 Overall Performance Ranking

Across five folds, the overall window-level ROC-AUC shows a clear performance ranking among the four configurations. The Temporal-Tile SegFormer baseline achieves the highest overall ROC-AUC (0.8659 ± 0.0213), followed by Temporal-Patchify SegFormer (0.8500 ± 0.0209) and MambaVision input256_512 (0.8477 ± 0.0231), while MambaVision input20 512 yields the lowest overall AUC (0.8192 ± 0.0246). This indicates that, under the same cross-validation protocol, SegFormer-based pipelines generally provide stronger discriminative capacity, whereas the MambaVision pipeline is more sensitive to the input mapping design.

#### 5.1.2 Effect of Input Encoding Size on MambaVision

Importantly, increasing the MambaVision input encoding from input20 512 to input256 512 yields consistent improvements across all folds. Using the per-fold mean AUCs, input256 512 outperforms input20 512 by +0.0310 (Fold 1), +0.0296 (Fold 2), +0.0229 (Fold 3), +0.0280 (Fold 4), and +0.0308 (Fold 5), resulting in an overall gain of +0.0285 AUC. The same trend is observed for the fold-wise best AUC values. These results suggest that a higher-capacity linear mapping layer with more channels can learn a more effective EEG channel mixing/feature projection prior to the vision backbone, thereby improving discriminability in a fold-consistent manner.

#### 5.1.3 Run-on-run Stability

In addition to absolute performance, we assess run-to-run stability via the within-fold standard deviation across six runs. Among the proposed configurations, Temporal-Patchify SegFormer exhibits the smallest variability, with fold-wise SDs ranging from 0.0033 to 0.0108, indicating superior stability and reproducibility across different random initialisations. In contrast, both MambaVision variants show larger fluctuations, particularly for input256 512 in Fold 3 (SD= 0.0312). Overall, these observations support the use of Temporal-Patchify Encoding as a robust input-feature mapping strategy and highlight that increasing the mapping capacity for MambaVision can substantially improve performance while still leaving room for further variance reduction.

### 5.2 Event-level performance

As discussed in the introduction and methods section, only event-level metrics can truly reflect the performance of a clinical seizure prediction system. To assess performance at this level, this research runs a pseudo-real-time prediction system on each event.

For each seizure event, both the preictal and interictal periods are segmented into 5-minute EEG signals, each comprising 150 2-second samples. By applying the EMA filter and then the K-out-of-N criteria, an alarm is triggered if K consecutive samples exceed the threshold.

Threshold values were selected from 0.5, 0.6, 0.7, 0.75, 0.8, 0.9. The resulting operating curve is shown in Fig. 6, where both sensitivity and false alarms per hour (FA/h) increase as the threshold decreases.

**Fig. 6.**
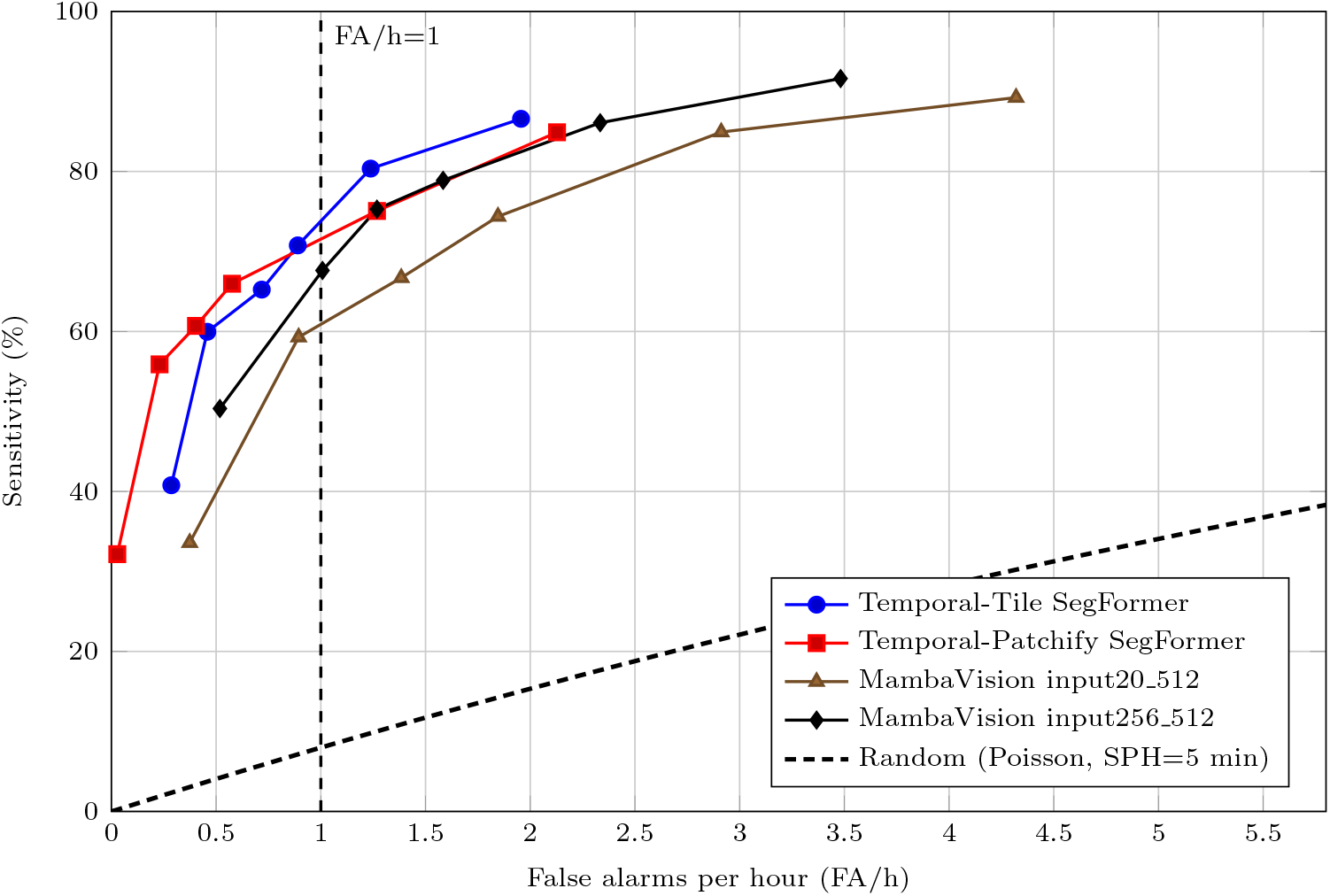
Event-level operating characteristics under EMA (*α* = 0.4) and persistence rule (*K* = 15).

#### 5.2.1 Performance at a Fixed Threshold

Table 6 sets a uniform threshold of 0.75 across four models and compares their performance at this operating point. At the same threshold, MambaVision input256 512 achieves the highest sensitivity 75.3%±11.3%, substantially higher than other models. Temporal-Patchify SegFormer yields the lowest false alarm rate (FA/h 0.40±0.28), substantially outperforming other models and demonstrating stronger false alarm control capabilities; although its sensitivity is relatively lower (60.7%±5.0%), it still remains stably above 60%. Temporal-Patchify SegFormer is suitable for deployment scenarios where the cost of false alarms is very high for patients.

**Table 6.**
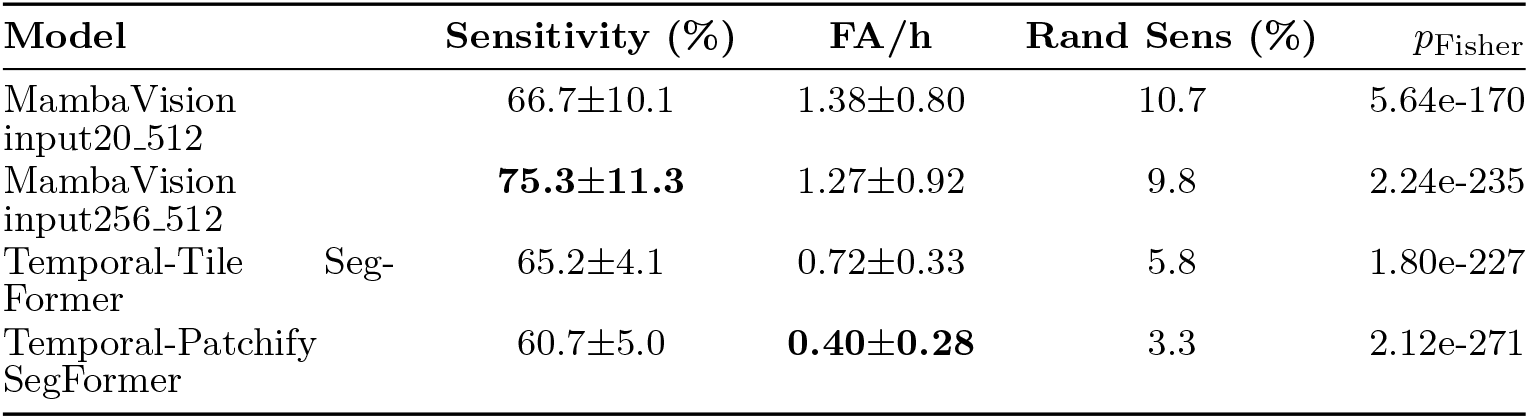
Operating point at threshold 0.75: event-level performance under EMA *α* = 0.4 and persistence *K* = 15. Values are mean±SD across 5 folds. “Rand Sens” is the expected sensitivity of a matched Poisson random predictor (SPH=5 min). *p*_Fisher_ combines five one-sided exact binomial tests across folds.

#### 5.2.2 Performance under a False-alarm Constraint

Table 7 shows that, under the uniform constraint FA*/*h *≤* 1, each model selects the threshold that maximises sensitivity while satisfying the false-alarm upper limit, enabling a fair comparison in the deployable regime. Under this constraint, Temporal-Tile SegFormer achieves the highest sensitivity within the controllable false-alarm range (Thr.=0.70, Sensitivity 70.8% ± 6.6%, FA*/*h = 0.89 ± 0.34), indicating the strongest prediction capability. Meanwhile, Temporal-Patchify SegFormer attains a more robust low-false-alarm operating point: at the same selected threshold (Thr.=0.70), it reduces the false-alarm rate to FA*/*h = 0.58 ± 0.35 (34.8% lower than Temporal-Tile) while maintaining a sensitivity of 66.0% ± 4.4%. In contrast, the MambaVision series requires higher thresholds to meet the low-false-alarm constraint (input20_512: Thr.=0.80; input256_512: Thr.=0.90), resulting in a larger drop in sensitivity. This suggests that MambaVision is better suited to operating regimes that tolerate higher false-alarm rates to achieve higher sensitivity, whereas maintaining high sensitivity under strict false-alarm constraints is more challenging.

**Table 7.**
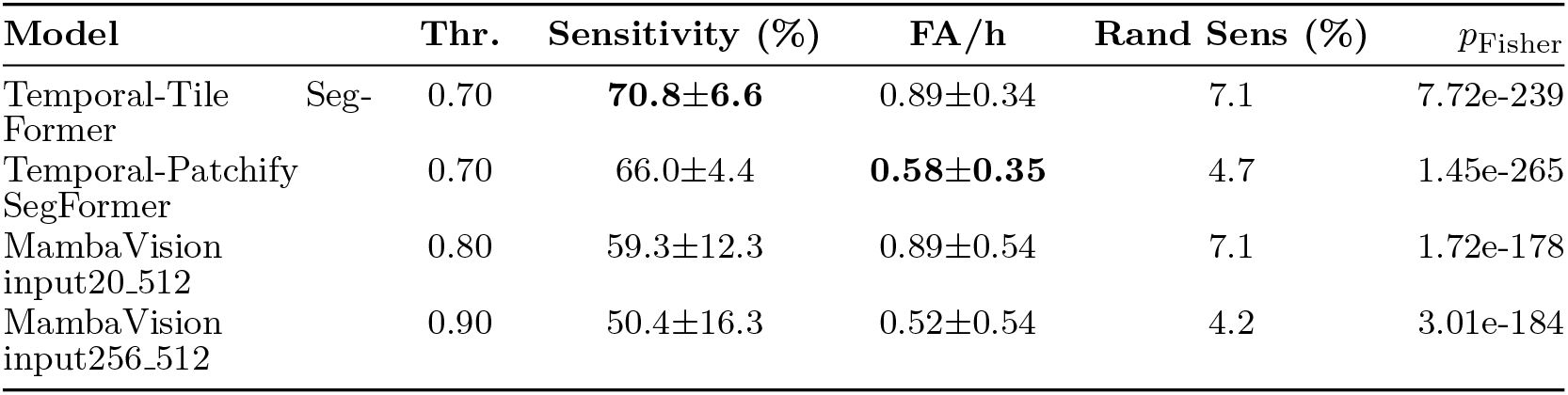
Best sensitivity under the deployable constraint FA/h *≤* 1 (EMA *α* = 0.4, *K* = 15). For each model, the threshold (Thr.) is selected to maximise sensitivity while satisfying FA/h *≤* 1. “Rand Sens” is the expected sensitivity of a matched Poisson random predictor (SPH=5 min). *p*_Fisher_ combines five one-sided exact binomial tests across folds.

Overall, SegFormer with proposed Temporal-Patchify excels at pushing the system into the “low false positive, deployable range.” In Table 6, Temporal-Patchify SegFormer has the lowest FA/h (0.40 ± 0.28), substantially lower than Temporal-Tile SegFormer (0.72 ± 0.33) and the two MambaVision models (1.27 ± 0.92, 1.38 ± 0.80), while maintaining a sensitivity of 60.7% ± 5.0%, which is considered a “sufficiently usable” prediction level. Table 7 further demonstrates that under the more realistic constraint of FA*/*h *≤* 1, Temporal-Patchify encoding can still suppress false positives to 0.58 ± 0.35 while maintaining a sensitivity of 66.0% ± 4.4%, demonstrating a more robust low-false-positive operating point.

Table 8 Partial AUC (pAUC) in the deployable region FA/h *≤* 1 (EMA *α* = 0.4, *K* = 15). pAUC is computed by truncating the event-level sensitivity–FA/h curve to FA/h *≤* 1. The Poisson random baseline assumes SPH = 5 min and has an analytic pAUC.

#### 5.2.3 Partial AUC Analysis

Moreover, Table 8 shows the partial AUC of each model. Since the operating point with *FA/h >* 1 on ROC is not deployable, partial AUC would be a meaningful metric for assessing each model’s performance. Our proposed Temporal-Patchify encoding method achieves the SOTA pAUC of 0.6094, 16.2% higher than the Temporal-Tile SegFormer baseline, again demonstrating the effectiveness of our proposed encoding method.

**Table 8.**
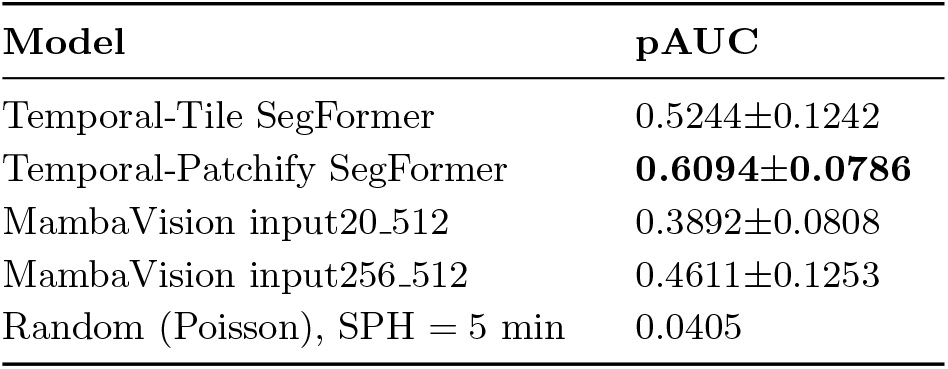
Partial AUC (pAUC) in the deployable region FA/h ≤ 1 (EMA α = 0.4, K = 15). pAUC is computed by truncating the event-level sensitivity–FA/h curve to FA/h ≤ 1. The Poisson random baseline assumes SPH = 5 min and has an analytic pAUC.

| Model | pAUC |
| --- | --- |
| Temporal-Tile SegFormer | 0.5244±0.1242 |
| Temporal-Patchify SegFormer | **0.6094±0.0786** |
| MambaVision input20_512 | 0.3892±0.0808 |
| MambaVision input256_512 | 0.4611±0.1253 |
| Random (Poisson), SPH = 5 min | 0.0405 |

### 5.3 Statistical validation

Fig. 6 shows the operating curve of a random Poisson predictor shown in equation 18, where *λ* is FA/h. It is clear that all 4 models outperform a random predictor.

To statistically validate event-level performance under matched false-alarm rates, we compared each operating point against a Poisson random predictor with the same FA/h (*λ*) and SPH = 5 min. The random hit probability is *p*_rand_ = 1 *− e*^*−λT*^ with *T* = 5*/*60 hour, yielding an expected random sensitivity of 100 *p*_rand_%. For each fold, significance is assessed via an exact one-sided binomial tail test on event hits, and fold-wise *p*-values are combined using Fisher’s method. We report statistical results at two pre-specified operating points: (A) a fixed threshold (Thr.=0.75) and (B) the best sensitivity under the deployable constraint FA*/*h *≤* 1.

Temporal-Patchify encoding achieves the lowest FA/h while maintaining clinically usable sensitivity (over 60%) and remains statistically significant compared with the matched random baseline.

### 5.4 Efficiency (inference throughput)

To assess each model’s capability for real-time deployment, inference throughput must be measured to determine whether it can function as a real-time system. We report inference throughput as the average number of EEG windows processed per second (windows/s).

All experiments were conducted on a Linux workstation equipped with an NVIDIA GeForce RTX 4090 GPU (23.54 GB VRAM), an Intel Xeon Gold 6336Y CPU @ 2.40 GHz (48 logical cores), and 251.3 GB system memory. Software versions were Python 3.11.14 and PyTorch 2.5.1 (CUDA 12.1, cuDNN 9.1.0), with NVIDIA driver 565.77.

For timing stability, we performed 1000 warm-up iterations followed by 10000 timed iterations. Throughput was computed from the total elapsed time covering the data-loader iteration, the host-to-device transfer, and the model forward pass (including the final activation), i.e., an end-to-end window-inference measurement (excluding disk I/O outside the loader). We additionally compute per-window latency (ms/window) using synchronised GPU timing around the forward pass; this excludes host-to-device transfer.

With non-overlapping 2-s windows, the theoretical real-time requirement is 0.5 windows/s. All configurations in Table 9 exceed this requirement by a large margin, indicating that inference speed is not a limiting factor in our setting; in practice, deployment performance is primarily determined by the chosen operating point on the sensitivity–FA/h curve. MambaVision input20_512 achieves the highest throughput, whereas expanding the mapping to input256_512 reduces throughput, consistent with increased computational cost. Within the SegFormer family, Temporal-Patchify is faster than Temporal-Tile, suggesting a more efficient construction of input features.

**Table 9.** Inference throughput measured as average windows processed per second (windows/s).

| Model | Windows/s |
| --- | --- |
| MambaVision input20_512 | **920.53** |
| MambaVision input256_512 | 510.76 |
| Temporal-Tile SegFormer | 610.70 |
| Temporal-Patchify SegFormer | 764.19 |

Nevertheless, the relative throughput differs across variants. MambaVision input20_512 achieves the highest throughput, whereas expanding the mapping to input256_512 substantially reduces throughput, consistent with the higher computational cost of the larger input projection. Within the SegFormer family, Temporal-Patchify is faster than the Temporal-Tile baseline, suggesting a more efficient input-feature mapping.

## 6 Conclusion

We propose a fully open and reproducible benchmark with a baseline for the scalp EEG seizure prediction community, serving as indicators for future reference. We report event-level key metrics including partial AUC and FA/h to assess models’ clinical usability. In detail. we converted the TUSZ dataset into an event-based 5-fold benchmark and replace the small-scale CHB-MIT dataset. SegFormers and the latest MambaVision are used for seizure prediction. We further compared all models with random predictors under the same false-alarm burden, showing that all evaluated models substantially outperformed the matched random predictors.

Our proposed Temporal-Patchify encoding achieves state-of-the-art performance over the Temporal-Tile SegFormer baseline [52], with a pAUC of 0.61, representing a 16.2% relative improvement under the same benchmark and evaluation protocol. This method enables the seizure prediction system to be deployed.

Overall, this work bridges the gap between seizure prediction algorithms and clinically usable seizure warning systems by providing an open, reproducible benchmark with event-level evaluation and deployment-oriented metrics.

Future work may focus on improving MambaVision’s performance by addressing its poor predictive capability through more suitable encoding methods. We expect this benchmark to serve as a reference protocol for future seizure prediction studies and to reduce over-optimistic reporting caused by sample-level evaluation.

## Declarations

### Ethics approval and consent to participate

This study performed a secondary analysis of publicly available, de-identified human EEG recordings from the Temple University Hospital EEG Seizure Corpus (TUSZ). The authors did not collect new data and had no interaction with human participants. Because only de-identified data were analyzed, additional ethical approval and informed consent were not required for this work.

### Consent for publication

Not applicable.

## Availability of data and materials

Access to the TUSZ dataset can be applied via [Neural Engineering Data Consortium](https://isip.piconepress.com/projects/nedc/html/tuh_eeg/). Code will be released upon acceptance.

## Competing interests

The authors declare that they have no competing interests.

## Funding

This work was supported by Basic Research Program BK20241815 of Jiangsu Province, Xi’an Jiaotong-Liverpool University Research Development Funding RDF-23-02-004, Jiangsu Industrial Technology Research Institute Postgraduate Research Scholarship FOS2402JAZ01 and funding A1503 from Apon Medical Ltd.

## Authors’ contributions

Ziyuan Yin: Conceptualisation, Data Collection, Methodology, Analysis. John Moraros: Supervision, Paper Revision. Shuihua Wang: Supervision, Methodology, Paper Revision.

## Acknowledgements

Acknowledgements to computing resources and medical knowledge guidance from Apon Medical Ltd.

## List of abbreviations

AUC: Area Under the Curve
CHB-MIT: Children’s Hospital Boston–MIT EEG Dataset
CNN: Convolutional Neural Network
EDF: European Data Format
EEG: Electroencephalography
EMA: Exponential Moving Average
FA/h: False Alarms per Hour
iEEG: Intracranial Electroencephalography
LSTM: Long Short-Term Memory
NEDC: Neural Engineering Data Consortium
pAUC: Partial Area Under the ROC Curve (low false-alarm region)
ROC: Receiver Operating Characteristic
ROC-AUC: Area Under the Receiver Operating Characteristic Curve
sEEG: Scalp Electroencephalography
SOP: Seizure Occurrence Period
SPH: Seizure Prediction Horizon
SOTA: State of the Art
STFT: Short-Time Fourier Transform
TIW: Time in Warning
TUSZ: Temple University Hospital Seizure Corpus
ViT: Vision Transformer

## Notes

https://isip.piconepress.com/projects/nedc/html/tuh_eeg/

### Competing Interest Statement

The authors have declared no competing interest.

